# Adaptation of Graph Convolutional Neural Networks and Graph Layer-wise Relevance Propagation to the Spektral library with application to gene expression data of Colorectal Cancer patients

**DOI:** 10.1101/2023.01.26.525010

**Authors:** Sebastian Lutz, Florian Auer, Dennis Hartmann, Hryhorii Chereda, Tim Beißbarth, Frank Kramer

## Abstract

**Motivation:** Colorectal Cancer has the second-highest mortality rate worldwide, which requires advanced diagnostics and individualized therapies to be developed. Information about the interactions between molecular entities provides valuable information to detect the responsible genes driving cancer progression. Graph Convolutional Neural Networks are able to utilize the prior knowledge provided by interaction networks and the Spektral library adds a performance increase in contrast to standard implementations. Furthermore, machine learning technology shows great potential to assist medical professionals through guided clinical decision support. However, the deep learning models are limited in their application in precision medicine due to their lack to explain the factors contributing to a prediction. Adaption of the Graph Layer-Wise Relevance Propagation methodology to graph-based deep learning models allows to attribute the learned outcome to single genes and determine their relevance. The resulting patient-specific subnetworks then can be used to identify potentially targetable genes.

**Results:** We present an implementation of Graph Convolutional Neural Networks using the Spektral library in combination with adapted functions for Graph Layer-Wise Relevance Propagation. Deep learning models were trained on a newly composed large gene expression dataset of Colorectal Cancer patients with different molecular interaction networks as prior knowledge: Protein-protein interactions from the Human Protein Reference Database and STRING, and pathways from the Reactome database. Our implementation performs comparably with the original implementation while reducing the computation time, especially for large networks. Further, the generated subnetworks are similar to those of the initial implementation and reveal possible, and even more distant, biomarkers and drug targets.

**Availability:** The implementation details and corresponding dataset including their visualizations can be found at https://github.com/frankkramer-lab/spektral-gcnn-glrp-on-crc-data

## 1 Introduction

Colorectal cancer (CRC) is one of the most prevalent cancer types in the world caused by gene alterations due to environmental factors or genetic inheritance. The four consensus molecular subtypes (CMS) identified for CRC indicate that highly variable molecular subnetworks differ not only between these sub-classes but also between single patients. Therefore, their gene expression profiles serve as a valuable base for patient stratification and personalized cancer prognosis. Those microarrays can be used as input features for classification tasks. The high-dimensionality of the gene expression data generates problems for its analysis, therefore, biological networks can help to reduce the complexity of possible interactions and introduce prior knowledge concerning the involved entities.

In recent years, improvements in machine learning approaches allow their application to enable advanced support in screening and diagnosis. In general, the models provide no insights into their decision-making process, raising legal and ethical questions, and consequently leading to a deprivation of acceptance in healthcare applications. Although most image-based analyses can be confirmed by medical experts, the application of deep learning models to different domains lacks professional human verification and validation. Therefore, various interpretation approaches were developed to be used in precision medicine to support the diagnosis and development of personalized treatment options (Yap *et al*., 2021; Chereda *et al*., 2021). Especially in cancer research, the large variability between different subtypes demands unique therapies for different patients (Rodriguez-Salas *et al*., 2017; Guinney *et al*., 2015).

Among the various deep learning models, Convolutional Neural Networks (CNN) already have proven their usefulness to gain prognostic insights in medical imaging for Alzheimer’s disease classification (Böhle *et al*., 2019) and breast cancer subtype prediction from previously analyzed gene expression data (Chereda *et al*., 2021; Bayerlová *et al*., 2017).

Layer-Wise Relevance Propagation (LRP) (Binder *et al*., 2016) is a framework to explain decisions from such classification tasks. There already exist LRP implementations for graph networks, for example, GNNExplainer or SubgraphX (Li *et al*., 2022) without supporting GCNNs. Thus, LRP was adapted as Graph Layer-Wise Relevance Propagation (GLRP) for the graph domain using the GCNN approach (Chereda *et al*., 2021). In general, the GLRP workflow looks as follows: In the first step, a patient’s data is mapped onto the graph vertices as input features representing an individual graph signal, and then classified by the GCNN model. Subsequently, the output classifications are propagated backward through the prediction network using the GLRP algorithm to calculate relevance scores for each input node. By applying a defined cut-off for the relevance score a patient-specific subnetwork can be constructed out of the remaining nodes.

Here, we present an adapted GCNN and GLRP approach using the recent graph neural network library Spektral (Grattarola and Alippi, 2021), and its application and validation on a novel comprehensive CRC dataset composed from multiple gene expression microarrays from the Gene Expression Omnibus (GEO) (Barrett and Edgar, 2006). We applied both models, the original GLRP implementation using GCNN and our adaptation using the Spektral library, with three different curated molecular networks as prior knowledge: Protein-protein interactions from the Human Protein Interaction Database (HPRD), binary interactions from the Reactome Pathway database, and functional protein association networks from the STRING database. Both implementations were then compared based on their performance and the generated patient-specific subnetworks to identify possible actionable genes.

## 2 Background and Materials

### 2.1 CRC dataset

The composed CRC dataset consists of multiple independent samples from the Gene Expression Omnibus (GEO) (Barrett and Edgar, 2006), an open online exchange platform and database for publicly available high-throughput gene expressions and hybridization arrays. Microarrays show transcriptional activity in each sample, which can be used to identify gene functions. Gene expressions describe the information of a gene generated by its transcription and translation process, which can be compiled into profiles to e.g. understand the genetic mechanisms of diseases or the response to complementary treatments proven to be helpful in cancer treatments (Brazma and Vilo, 2000; Slonim and Yanai, 2009).

Each microarray was extracted using the Affymetrix HG-U133 Plus 2.0 and is represented as *ExpressionSets* using the Biobase R-package (R. Gentleman, V. Carey, M. Morgan, S. Falcon, 2017). The dataset contains 733 patients having the following GEO *Series* accession codes: GSE15960, GSE22598, GSE4183, GSE33113, GSE18105, GSE73360, GSE20916, GSE4107, GSE24514.

Every patient was binary labeled by their current status (cancerous or not cancerous), resulting in 303 healthy and 434 cancerous samples. Preprocessing was done automatically by first applying the Robust Multi-Array Average Algorithm (RMA) (Irizarry *et al*., 2003) on each *Series* by background correcting, normalizing, and summarizing the data. Next, the used microarrays were combined, based on the probe ID mappings of the corresponding Affymetrix mappings (hgu133plus2.db, 2022), and quantile normalized over the whole set. Finally, the CRC subtypes were predicted for each patient sample, using the CMSCaller R-package (Eide *et al*., 2017), which is based on the Nearest Template Prediction (NTP) algorithm (Hoshida, 2010).

### 2.2 Prior knowledge from biological networks

Three commonly used molecular interaction networks were used as prior knowledge to model the complex interactions between genes. The single networks thereby were chosen to differ in their size to properly estimate this impact on model performance. The networks were automatically obtained from the Network Data Exchange (NDEx) (Pratt *et al*., 2015), an open-source online platform for editing, sharing, and distributing networks. NDEx uses the Cytoscape exchange (CX) data format, inspired by the Cytoscape (Franz *et al*., 2016) open-source visualization software to store graph data as separated aspects of the network. It also allows for custom metadata fields that are added to the resulting subnetworks to not only publish them onto NDEx but create visualizations using the MetaRel-SubNetVis web application (Auer *et al*., 2022).

The first used network is the Human Protein Reference Database (HPRD, Version 9 from 2009) (Keshava Prasad *et al*., 2009) containing manually curated and experimentally proven protein-protein interactions. containing manually curated and experimentally proven protein-protein interactions. After mapping the network to the gene expression data the resulting graph contains 6888 vertices and 27841 edges and is available on the NDEx platform (UUID: *079f4c66-3b77-11ec-b3be-0ac135e8bacf*). Reactome is an openly available online platform for peer-reviewed and curated human physical and pathological pathways and reactions (Gillespie *et al*., 2022). A merged, binary network of the Reactome database with its nomenclature converted to Human Genome Organization (HUGO) gene symbols (Seal *et al*., 2023) resulting in 7926 vertices and 194502 edges was also obtained from NDEx (UUID: *4bc71515-86da-11e7-a10d-0ac135e8bacf*).

As a third molecular network, an interaction network from the Search Tool for the Retrieval of Interacting Genes (String) is used. This database features not only proven but computationally predicted protein-protein physical interactions and functional associations (Szklarczyk *et al*., 2021). The graph contains 17185 vertices and 420534 edges (UUID: *275bd84e-3d18-11e8-a935-0ac135e8bacf*).

### 2.3 Deep learning model

Machine learning models can produce meaningful outputs from their given inputs to retrieve rules generalizable to unseen data. This is achieved by using the predicted output to measure its deviation from the desired output and adjust the chosen learning algorithm accordingly. Here, the neural network is trained for the classification of patients into cancer subtypes using their gene expression data as input features representing graph signals. Convolutional Neural Networks Convolutional Neural Networks (CNN) are able to capture the spatial locality of neighboring pixels based on the idea, that key patterns in grid-like data structures like pictures make up only a small section of it (Goodfellow *et al*., 2016). A simple CNN consists of at least one convolutional layer, followed by a pooling layer to downsample the input further, and a fully connected neural network to classify the downsampled input.

#### Graph Convolutional Neural Networks

Since CNNs can’t work with arbitrary data structures in the non-Euclidean domain, GCNNs were developed to capture the correlation of neighboring nodes in graph networks, being able to classify network-based input. Therefore, a different propagation rule is needed compared to a normal convolution. In this case, the GCNN uses Chebyshev Polynomials (ChebNet) to approximate the spectral graph convolutions (Defferrard *et al*., 2016).

#### Graph Layer-wise Relevance Propagation

Since interpretability is a requirement for neural networks used in precision medicine, LRP (Binder *et al*., 2016) was developed as a technique to explain the decisions of a CNN by propagating the predictions with their corresponding weights and activations backward through the network. Multiple LRP strategies exist to adapt the propagation to the model architecture. After calculating all relevance scores, they can be mapped on the input pixels creating a heatmap thus illustrating the importance of each pixel for the networks’ decision. The GLRP adapts the LRP to GCNNs as a post-calculated step. Since GCNNs work on graphs, the relevance score now represents the importance of each node, and therefore each gene, for the prediction of the cancer sub-type. Setting a threshold for the gene relevance score allows spanning a patient-specific subnetwork indicating alterations in the genetic interactions relevant for cancer progression and patient treatment.

## 3 Methods

### 3.1 Pre-Processing

This implementation separates the steps to retrieve the data, their pre-processing, training of the model, and post-processing to generate the corresponding results to provide modularity to the setup. Since reproducibility is of major concern, every step is automatized and configurable via extensive configuration files managing model hyperparameters, data paths, or visualization meta data. In the pre-processing step, the raw data from the GEO database is transformed into Spektral readable data structures.

At first, the GEO accession codes for the corresponding Series are used to automatically request, unpack, and clean the raw data in form of Affymetrix CEL files from the GEO database. The RMA algorithm is applied to the raw microarray data and normalized using the Affy R-package (Gautier *et al*., 2004). The *annotate* R-package (R Gentleman, 2022) annotates the data with the corresponding HUGO gene names.

After each file is processed individually they are merged into one data frame and cleaned by removing missing values. Finally, the dataset is quantile normalized. The CRC subclasses for each sample are predicted using the CMSCaller R-package (Eide *et al*., 2017). Finally, the combined gene expression dataset must be mapped to the configured molecular network based on their gene ID to assign to each node its corresponding gene expression value.

Spektral uses its own data model to represent input graphs and features, therefore, the labels, feature values, and molecular network are converted into Spektral Graphs. A Spectral Graph represents the data for one patient as an adjacency matrix of the molecular network, corresponding nodes and edge features, and the corresponding ground truth. The single Spektral graphs are coarsened and put into a combined Spektral DataSet, using built-in manipulation functions.

Spektral features multiple data loaders for training supporting different modes. We use the MixedLoader since only one adjacency matrix for all input features is required when using graph signals. Hence, the adjacency matrix is loaded only once for each sample, leading to reduced memory consumption. Nevertheless, it does not support internal helper functions and therefore requires a custom training loop.

### 3.2 Spektral model

This implementation uses Spektral (Grattarola and Alippi, 2021), an opensource library supporting graph neural networks implemented with TensorFlow (TensorFlow Developers, 2022). In contrast to the original implementation, Spektral uses TensorFlow 2 instead of TensorFlow 1.15. The required update to the new version introduces eager execution and simplifies the model-building workflow with the inclusion of the Keras framework as the high-level API. Therefore, it allows for faster debugging, development, and deployment. Furthermore, TensorFlow 2 endorses more fine-grained control of subclassed models and custom training loops. Although more effort is required for the creation of neural network stages this results in potential performance gains when optimizing each stage individually. Spektral implements custom TensorFlow Layers, which provide functions for further customizable convolutional layers since certain layers require additional pre-processing on their inputs. For example, it already implements a working ChebNet convolutional layer, with an exception for clustering the input graphs as used in CNNs with fast localized spectral filtering.

As shown in **Fig. 1** the created model consists of two convolutional layers with their corresponding Max- or Average-Pooling layers, followed by a Flatten layer. Lastly, the convolved data is fed through two fully connected layers and one output layer with a SoftMax activation function. Every other convolutional and fully connected layer uses the Rectified Linear Unit (ReLU) activation function. Layered coarsened graph inputs are passed into the network using two input layers for each convolution, while the adjacency matrix is passed via a third input layer. The latter is reused since it does not change throughout the network.

**Fig. 1:**
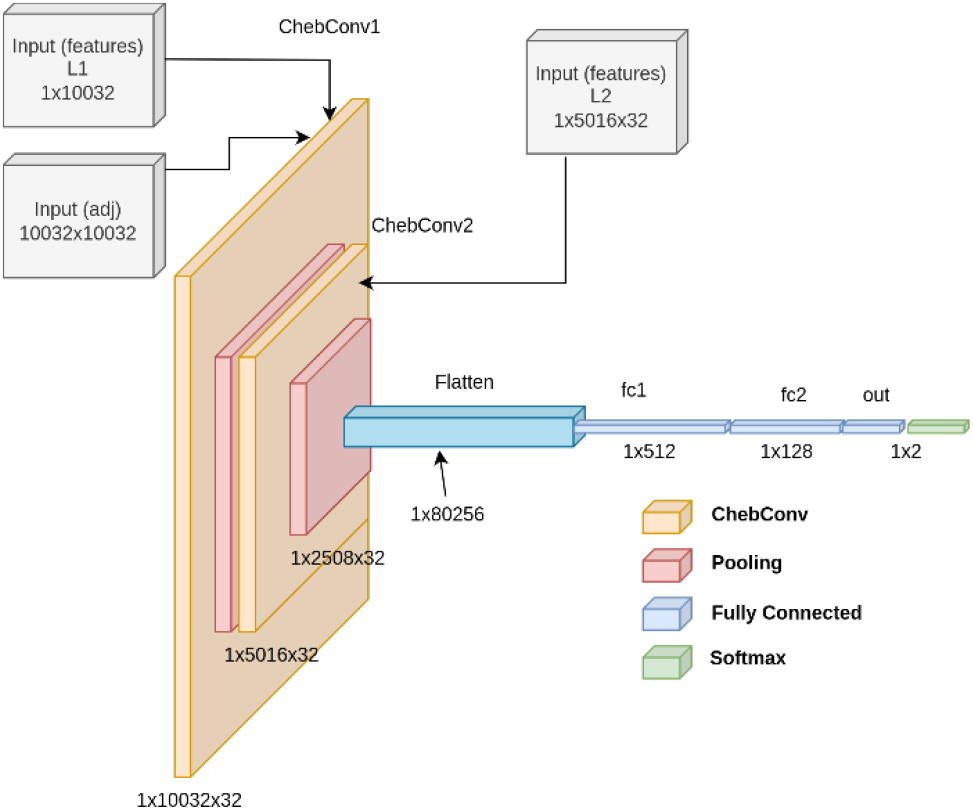
Neural network architecture of the used GCNN for the HPRD PPI network with their given input sizes after pooling the graphs.

Instead of the sequential or functional Keras API, a subclassed model was used due to the required intermediate input layers for the coarsened graph inputs in conjunction with the custom training loop. It implements a Keras Model inside the Model module and overwrites the call function feeding inputs through the network. Furthermore, subclassing allows adding custom functions to use on the network itself but limits the use of higher-level TensorFlow features. Since the model structure and its corresponding weights must be retained for the GLRP in the post-processing, the model implements a function to return a built Keras graph with the corresponding correct input sizes. Also, callbacks and higher-level TensorFlow metrics are not supported, therefore lower-level metric updates were re-implemented. The metrics Accuracy, Area Under Curve (AUC), Precision, Recall, and the F1-Score were used to reflect the model performance on the training and test set.

The datasets were separated into batches by the MixedLoader and fed through the custom training loop, using 10-fold cross-validation on the training set and a test set including ten percent of the samples.

### 3.3 Post-Processing

Post-processing was performed using the trained model to propagate the outputs back through the network. The relevance scores were calculated using GLRP and subsequently, the patient-specific subnetworks and visualizations were generated. Afterward, the output graphs were converted to the CX format and finally uploaded to the NDEx platform.

The GCNNExplainer module uses the test dataset to calculate the relevance scores by applying different propagation rules on different layer levels as stated previously. Thus, all layers are divided into three groups, excluding those without non-trainable weights. Those groups then are each assigned to the basic rule for the lower layers, the ϵ-rule for the middle layers, and the γ-rule for the upper layers. Since positive relevances should only exist, the z^+^-rule is applied additionally to all layers by clipping the weights beforehand. A generic function calculates the ρaccordingly.

Polynomials of the coarsened adjacency matrices are calculated before propagating them through the network to save computing time by reversing the layers from the loaded model and looping through them. Since activations, in contrast to the weights, of each preceding layer are not saved within the model a slice up to the currently used layer is converted to a Keras Model to retrieve the intermediate activations. They then are passed with the weights and if necessary pre-calculated polynomials to the corresponding propagation function. A propagation function uses the activations, weights, the previous relevance score, and the given rule to return the relevances before the current layer. When using ChebNet layers, the relevances have to be propagated through each output filter using the corresponding Chebyshev polynomial calculated before (Chereda *et al*., 2021).

Finally, the combined relevance scores can be used for the visualization. A threshold for the relevance scores is defined on a patient basis to span patient-specific subnetworks from the 140 most relevant genes. The resulting network has to comply with the CX data model from NDEx to work with the visualization platform MetaRelSubNetVis. Therefore, additional metadata properties are required in the converted CX network to determine network-specific visualization parameters for each sample. The resulting graph is built by using the molecular network as a base and compiling the needed properties i.e., the quantile levels and the standard deviation from the resulting graphs. Those converted networks are then uploaded to NDEx using the NDEx2 python client.

### 3.4 Data and source code availability

The source code (GPL-3.0 License) and all necessary data to reproduce the results can be found at https://github.com/frankkramer-lab/spektral-gcnn-glrp-on-crc-data. The repository includes the Spektral framework itself, the generated results for the CRC dataset, and the GCNN and GLRP Spektral implementation. Additionally, it features documentation of the results including the corresponding visualizations and code. Furthermore, the included command-line tool can be used to generate visualizations or train models based on the configuration files.

## 4 Results

To compare both, the original and our Spektral implementation, all results were generated on the same workstation with an Nvidia RTX 3070, AMD R3700X, and 32GB of RAM. The patient-specific networks of the CRC dataset are combined into one for each model can be found on the NDEx platform. They can be interactively investigated using the MetaRelSubNetVis web application, which can load the networks directly from the NDEx platform.

The performance of the different models consists not only of the models’ prediction but also training and GLRP calculation times. The computing time took 937 up to 8,838 seconds on average, when using the Spektral implementation, in contrast to the original implementation, which took 967 to 16,536 seconds to complete depending on the used network. In **Table *1*** also the corresponding scores for the CRC dataset are displayed. Each model achieves >90% accuracy in each implementation and has comparable performance scores in general: the accuracy stays about the same, but the Spektral implementation shows an improvement in accurracy and F1-Score of one to two percentage points, even though the AUC score is two to four percent less.

**Table 1.** Performance of the original and Spektral implementations

| Models | Accuracy | F1-Score | AUC*100 | Avg. Time in s |
| --- | --- | --- | --- | --- |
| *HPRD PPI (6888, 27841)* |  |  |  |  |
| Original GCNN | 94.40 | 94.50 | **98.50** | 967 |
| Spektral GCNN | **95.95** | **96.96** | 95.92 | **937.4** |
| *Reactome (7926, 194502)* |  |  |  |  |
| Original GCNN | 93.20 | 93.20 | **98.80** | 4040.4 |
| Spektral GCNN | **95.95** | **95.96** | 96.92 | **1231.3** |
| *String (17185, 420534)* |  |  |  |  |
| Original GCNN | 93.20 | 93.20 | **97.50** | 16536 |
| Spektral GCNN | **93.24** | **94.85** | 93.92 | **8838** |

Performance of the original implementation and Spektral implementation on the CRC dataset with 733 samples, using the three different networks with their corresponding number of vertices and edges in brackets respectively.

The ten topmost occurring genes of each implementation are listed in **Table *2*** and sorted by the corresponding used network. Genes occurring in multiple implementations are marked in orange when appearing in one and green when appearing in both of them. It is shown that the most selected genes are mostly similar to the original implementation using the smallest network (HPRD). When using larger networks as prior knowledge, such as the Reactome pathways and STRING PPI network, the overlap tends to decrease.

**Table 2.**
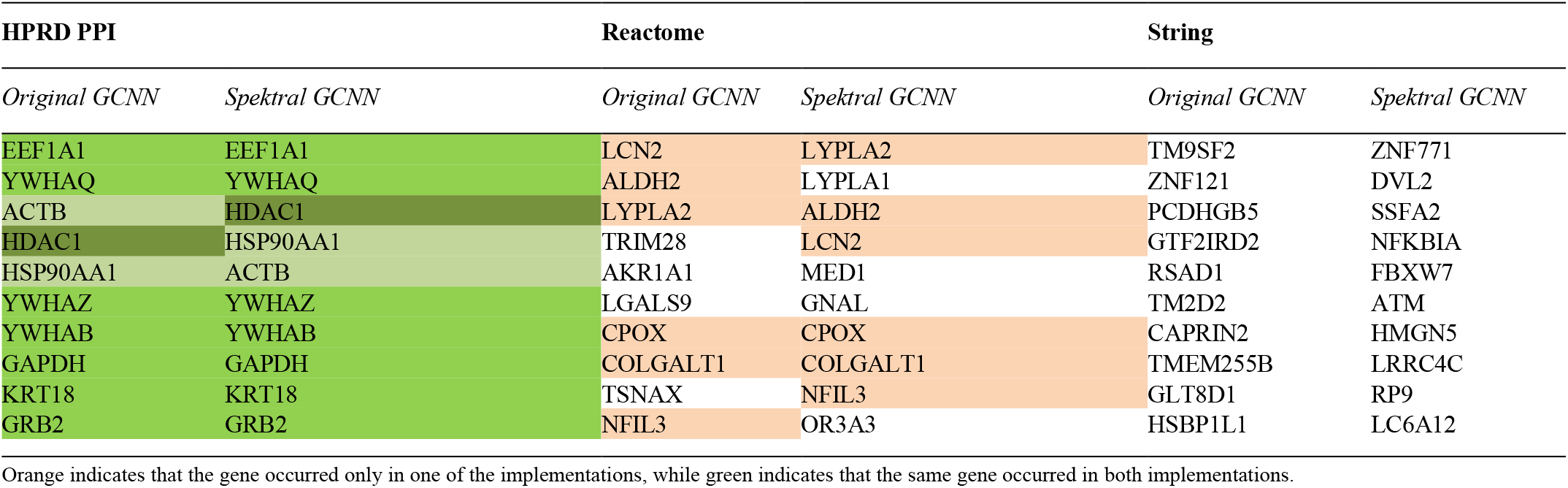
Most occurring genes for the CRC dataset for each used network compared by each implementation.

Two representative samples (**Table *3***) were selected to illustrate the differences between cancer and healthy samples. The corresponding sub-networks for each implementation method and network are examined and visualized by gene expression level using the MetaRelSubNetVis web application visualize (**Fig. *2***). The gene expression levels are color coded with red representing a high level (>75% quantile), yellow a medium level (<75% and > 25% quantile), and blue a low level (<25% quantile) of gene expression. Both implementations generated mostly similar subnetworks for each sample according to their contained genes, their occurrences, and the gene expression levels.

**Table 3.** Sample patients

| Patient | Subtype | Classification |
| --- | --- | --- |
| GSM93946 | Normal | No-Cancer |
| GSM452660 | CMS1 | Cancer |

Used patient samples to visualize them with their corresponding subtypes and labels

**Fig. 2.**
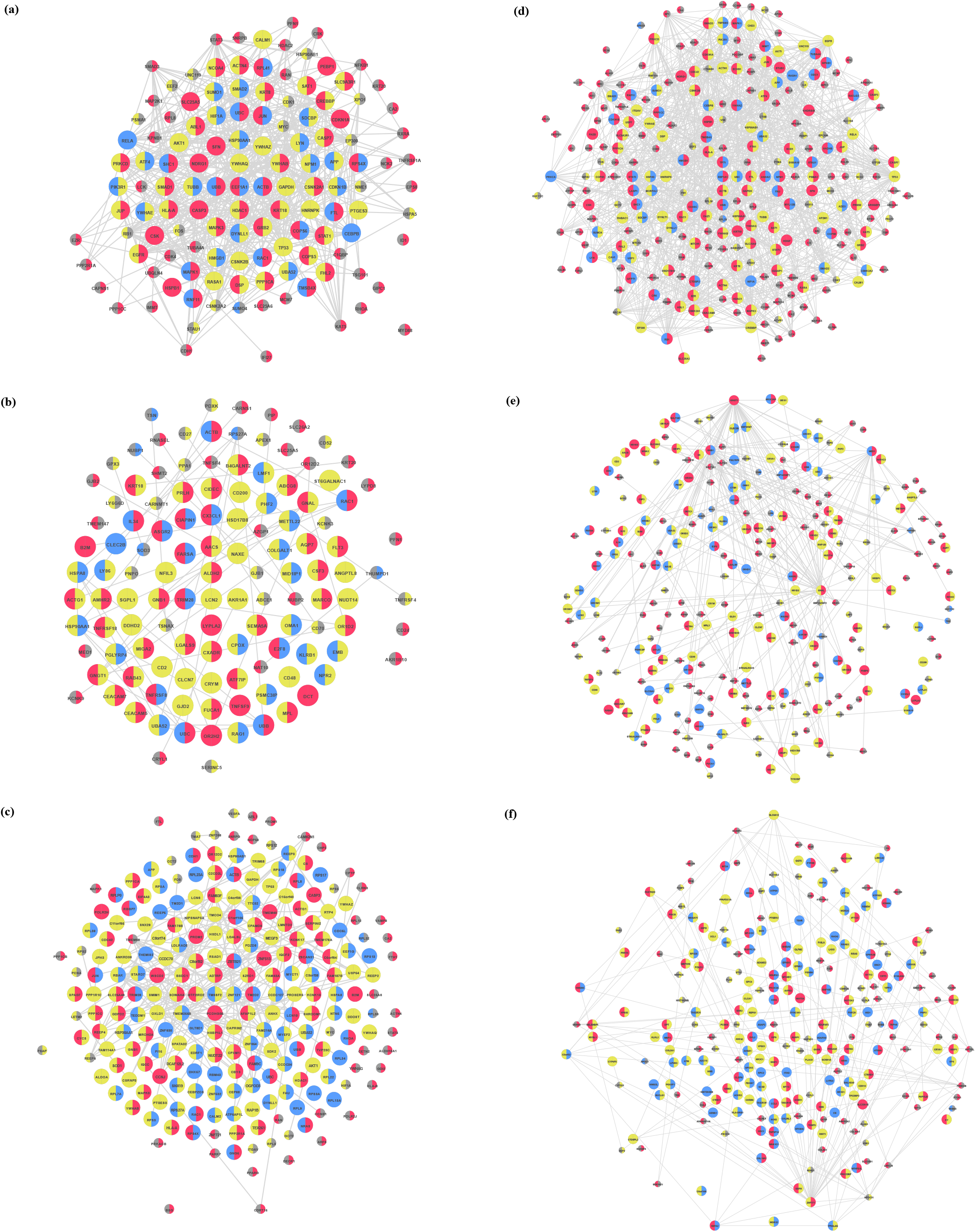
Comparison of two sample subnetworks from the CRC dataset for the patients GSM 93946 (No-Cancer) and GSM 452660 (Cancer), using the original (left, a-c) and Spektral implementation (right, d-f) respectively. The rows show visualizations of the subnetworks based on the different source networks from HPRD, the Reactome pathway database, and the String database.

## 4 Discussion

The Spektral implementation is evaluated based on the performance scores and the resulting subnetworks including the three different used molecular networks. Our implementation shows improved computation times compared to the original implementation, especially with increasing network size. However, when splitting training and GRLP computation, the original implementation is comparably faster in training. This is primarily due to the improved data handling from the Spektral model by retaining the adjacency matrix for each batch sample, as well as low-level improvements from TensorFlow 2 itself. An explanation for the slower GLRP computation is the fact, that the original implementation keeps each activation for each layer in the RAM. Hence, they do not have to be calculated for each layer again.

Another problem regarding the original implementation was the inability to save the model before computing the GLRP, regardless of the saved checkpoints. Therefore, the model had to be trained every time and could not be re-used with e.g., different datasets independently, which was fixed using modular independent steps. Since it is not easily possible to save intermediate activations with Spektral and TensorFlow layers, the GLRP for Spektral compiles a slice of the original model and propagates the inputs through it to recompute the intermediate activations. This trade-off ensures the execution of the GLRP for any given model without depending on temporarily stored information from previous steps.

Our implementation shows higher accuracy on average and better F1-Scores on the dataset, yet the original implementation yields a comparable AUC result. This might be a phenomenon due to class imbalance but could possibly indicate overfitting issues. The only significant difference from the standard implementation is using Spektral-specific layers and current versions of TensorFlow with its corresponding libraries. Hence, performance improvement is primarily based on different low-level improvements. Further improvements could be made using an improved convolutional layer trying to fix problems due to illegally learned coefficients within the original version. This leads to a possible fix for overfitting issues and the Runge phenomenon (He *et al*., 2022).

Not only performance plays an integral part in measuring the abilities of the new implementation: the generated subnetworks must be meaningful in correspondence to the original implementation. Within the single molecular networks, both models show great overlap in the most relevant genes for their prediction. Across the different networks, it can be seen that the smaller datasets still perform well, and the topmost occurred genes are overlapping. With the STRING network being larger than both other networks, this circumstance might help to uncover more distant associations. An obvious reason is the increased number of edges and genes since the table shows only the most occurring ones. Thus, fewer occurring genes would be hidden in resulting gene list from **Table 2**, although being relevant for specific cancer sub-types or individual patients. Regarding the occurred genes in the CRC dataset and the HPRD PPI network, the most occurred genes e.g. EEF1A1, YWHAQ, YWHAZ, YWHAB, or ACTB are already associated with a variety of cancer types and are already used for diagnostics and therapies (Guo *et al*., 2013; Kim *et al*., 2019; Gan *et al*., 2020).

Since the original and Spektral implementations did not yield many different results, it underlines the assumed correctness of the results from both Spektral layers.

The ALDH2, LYPLA2, LCN2, and CPOX genes occurring as results of the Reactome pathways are associated with cancer too: For example, ALDH2 is known for alcohol degradation, but also an indicator of tumor development. LCN2 is directly linked to colorectal cancer development (Zhang and Fu, 2021; Crous-Bou *et al*., 2013; Pasche *et al*., 2002; Fitzgerald *et al*., 2018; Srinivasalu *et al*., 2020; Feng *et al*., 2016).

The computed topmost occurrences using the STRING network show, that selected genes, e.g. CAPRIN2 or TM9SF2, are linked to the survival of colorectal cancer patients (He *et al*., 2021; Clark *et al*., 2018). The Spektral implementations also selected multiple genes related to colorectal cancer, e.g. the DVL2, SSFA2, NFKBIA, and FBXW7. FBXW7 is used in colorectal cancer prognosis, while NFKBIA is associated with the risk of CRC. Those findings again support the hypothesis that the STRING network is able to uncover more distant associations.

The corresponding generated subnetworks for the CRC dataset (**Fig. *2***) reflect the gene expression levels according to the occurred genes. For example, ACTB or YWHAB using the HPRD PPI or SELENBP1 using the STRING network are more expressed in CRC samples compared to non-cancer ones.

## 6 Conclusion

In this work, we introduce a performant adaption of GCNN and GLRP using the Spektral framework for the attribution of prognostic relevance to the single genes of a biological network. Furthermore, a new dataset with CRC samples was assembled and validated using three different molecular networks as prior knowledge, and compared to the previous implementation.

The resulting subnetworks of the CRC dataset using our method show high similarity to the original implementation and only slightly differ regarding their gene occurrences and corresponding associations providing valid marker genes for colorectal cancer progression. Moreover, our implementation requires less computation time, especially when using large molecular networks as prior knowledge. The predictive performance is comparable if not better in several tasks than the original implementation. Taking the modularity of each pipeline step into account, our Spektral-based approach is much more user-friendly, while being able to produce comprehensible results compared to the original implementation.

We were capable to identify genes already known to be associated with colorectal cancer risk and progression, proving the effectiveness of our model for diagnostic purposes. In addition, the many detected genes were overlapping between both implementations, and even across networks used as prior knowledge, indicating fundamental biological connections. Since also the larger networks detect common genes although they feature more genes, they potentially can uncover more distant patient- or subtype-specific alterations. Further, our model could generalize reasonably well, even with an unbalanced and small dataset, and therefore should be able to produce relevant results, even for previously unseen data.

